# Fuzzy Quantification of Common and Rare Species in Ecological Communities (FuzzyQ)

**DOI:** 10.1101/2020.08.12.247502

**Authors:** Juan A. Balbuena, Clara Montlleó, Cristina Llopis-Belenguer, Isabel Blasco-Costa, Volodimir L. Sarabeev, Serge Morand

## Abstract

1. Most species in ecological communities are rare whereas only a few are common. This distributional paradox has intrigued ecologists for decades but the interpretation of species abundance distributions remains elusive.

2. We present Fuzzy Quantification of Common and Rare Species in Ecological Communities (FuzzyQ) as an R package. FuzzyQ shifts the focus from the prevailing species-categorization approach to develop a quantitative framework that seeks to place each species along a rare-commonness gradient. Given a community surveyed over a number of sites, quadrats, or any other convenient sampling unit, FuzzyQ uses a fuzzy clustering algorithm that estimates a probability for each species to be common or rare based on abundance-occupancy information. Such as probability can be interpreted as a commonness index ranging from 0 to 1. FuzzyQ also provides community-level metrics about the coherence of the allocation of species into the common and rare clusters that are informative of the nature of the community under study.

3. The functionality of FuzzyQ is shown with two real datasets. We demonstrate how FuzzyQ can effectively be used to monitor and model spatio-temporal changes in species commonness, and assess the impact of species introductions on ecological communities. We also show that the approach works satisfactorily with a wide range of communities varying in species richness, dispersion and abundance currencies.

4. FuzzyQ produces ecological indicators easy to measure and interpret that can give both clear, actionable insights into the nature of ecological communities and provides a powerful way to monitor environmental change on ecosystems. Comparison among communities is greatly facilitated by the fact that the method is relatively independent of the number of sites or sampling units considered. Thus, we consider FuzzyQ as a potentially valuable analytical tool in community ecology and conservation biology.

## Introduction

Ecological communities are formed by species that differ widely in abundance. Almost invariably the observation is that most species are rare, whereas a few are common (Magurran & Henderson, 2011). This pervasive pattern has intrigued ecologists for decades but, despite the large literature on the topic, the interpretation of species abundance distributions remains elusive (Werner et al., 2014; Enquist et al., 2019). The assumption often made is that underlying factors, such as immigration, succession and competition, eventually determine differences in establishment and persistence of each species in the community (McGill et al., 2007; McGill, 2011; Alroy, 2015; Calatayud et al., 2019).

A quantitative framework for species commonness and rarity amenable to hypothesis testing and statistical modelling would facilitate evaluating the roles played by demographic variables and species traits, thereby illuminating assembly rules in ecological communities. Such a framework would also be extremely valuable for conservation biology in at least three important aspects: (1) Species rarity is an important predictor of extinction risk because the impact of environmental disturbances is expected to be higher on small populations (Davies, Margules, & Lawrence, 2000) and since budgets for biodiversity conservation are limited, quantifying the rarity of species would facilitate prioritizing some over the others. (2) From a functional perspective, the role played by common and rare species in providing ecosystem services is currently under scrutiny. Although intuitively it might be assumed that most of the ecosystem functionality should reside in the former, the contribution of rare species is still poorly understood and, in fact, might be substantial (Leitão et al., 2016; Violle et al., 2017; Dee et al., 2019). So assessing rarity could also be justified in terms of identifying species that provide essential ecosystem services (Flather & Sieg, 2007; Violle et al., 2017; Dee et al., 2019) or stabilize ecological communities (Calatayud et al., 2019). (3) Monitoring variation of commonness-rarity patterns over time or along geographical and environmental gradients provides a simple way to obtain crucial information on ecosystem changes (McGill, 2011). For instance, if common species become increasingly rare in response to environmental disturbances, it might have a cascading effect on the rest of the community (Gaston & Fuller, 2008).

Thus, metrics of commonness and rarity at species and community level would be extremely useful to unveil the architecture of ecological communities, assess the likelihood of extinction of rare species, correlate commonness or rarity with functional distinctiveness and monitor environmental change. However, a universal quantitative framework is currently lacking. A great deal of effort has been put on establishing the distribution patterns emerging from the categorization of species as common or rare (Gray, Bjørgesæter, & Ugland, 2005; McGill et al., 2007; Antão, Connolly, Magurran, Soares, & Dornelas, 2017). However, a major problem of fitting models to species abundance distributions has been adjusting the data to a suitable theoretical distribution (Williamson & Gaston, 2005; McGill et al., 2007; Alroy, 2015). To some extent this is because the border between common and rare species is often blurred (Magurran & Henderson, 2011), which has led authors to propose additional subcategories of rarity (Hanski, 1991; Yu & Dobson, 2000; Arnan, Gaucherel, & Andersen, 2011).

Herein we propose shifting the focus from species categorization to a quantitative approach that seeks to place each species along a rare-commonness gradient. Fuzzy Quantification of Common and Rare Species in Ecological Communities (FuzzyQ) is based on the analysis of abundance-occupancy relationships (AORs), which assumes a positive relationship between local abundance and occupancy (Gaston et al., 2000; Gaston & He, 2011). Given a community surveyed over a number of sites, quadrats, or any other convenient sampling unit, FuzzyQ applies a fuzzy clustering algorithm (Kaufman & Rousseeuw, 1990) that estimates a probability for each species to be common or rare based on its AOR.

Although widely used, we acknowledge at the onset that abundance and/or occupancy are not the only criteria to assess species commonness and rarity (Gaston, 1994, 1997). However, the key point is that regardless of the data used, we can always use fuzzy clustering to quantify the degree of belonging of each species to the common or rare categories (or any other pre-established categorization for that matter).

We show herein that FuzzyQ produces ecological indicators easy to measure and interpret that are amenable to hypothesis testing and statistical modelling. In addition, FuzzyQ is distribution free, i.e. no a priori assumption about the distribution of species abundances is required. We illustrate the capabilities of the framework with two real-world examples involving each related and unrelated (i.e. not sharing species) communities and evaluate the effect of sample size on the estimation of commonness and rarity.

## Overview of FuzzyQ

FuzzyQ is provided as an R package (R Core Team, 2020), available at https://github.com/Ligophorus/FuzzyQ, which depends on algorithms implemented in package cluster (Maechler, Rousseeuw, Struyf, Hubert, & Hornik, 2019). We first illustrate application of FuzzyQ with a dataset of ant species (ants_Darwin_A in Calatayud el al., 2019) comprising the abundance of 46 species in 100, 18×18 m plots sampled in the Northern Territory, Australia (Arnan et al., 2011).

Table 1 provides an overview of the functions in package FuzzyQ. Function `fuzzyq` takes a given site-by-species abundance matrix and performs a fuzzy clustering algorithm that evaluates all pairwise dissimilarities among species in terms of their AORs to allocate each species into two clusters of common and rare species, respectively. Since occupancy and abundance are in different scales and can come in different units (for instance, the former can be reported as either number or fraction of sites occupied), `fuzzyq` uses by default Gower’s (1971) dissimilarities, which are appropriate for such mixed data. Clustering is subsequently performed with function `fanny` in `cluster`, which aims to minimize the objective function

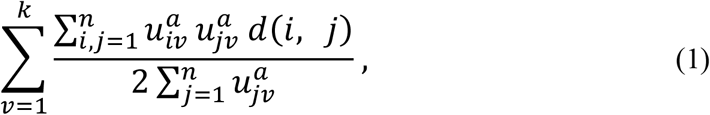

where *u*_*iv*_ and *u*_*jv*_ are the membership coefficients of observations (species in our case) *i* and *j* to cluster *v*, *n* is the number of observations, *k* is the number of clusters (herein 2: common and rare), *a* is a membership exponent (we set *a* = 2 as in the original formulation of Kaufman & Rousseeuw 1990) and *d*(*i*, *j*) is the dissimilarity between observations *i* and *j* (Maechler et al., 2019). Fig. 1a displays the `fuzzyq` allocation of species of the ant dataset to the rare and common clusters.

**FIGURE 1.**
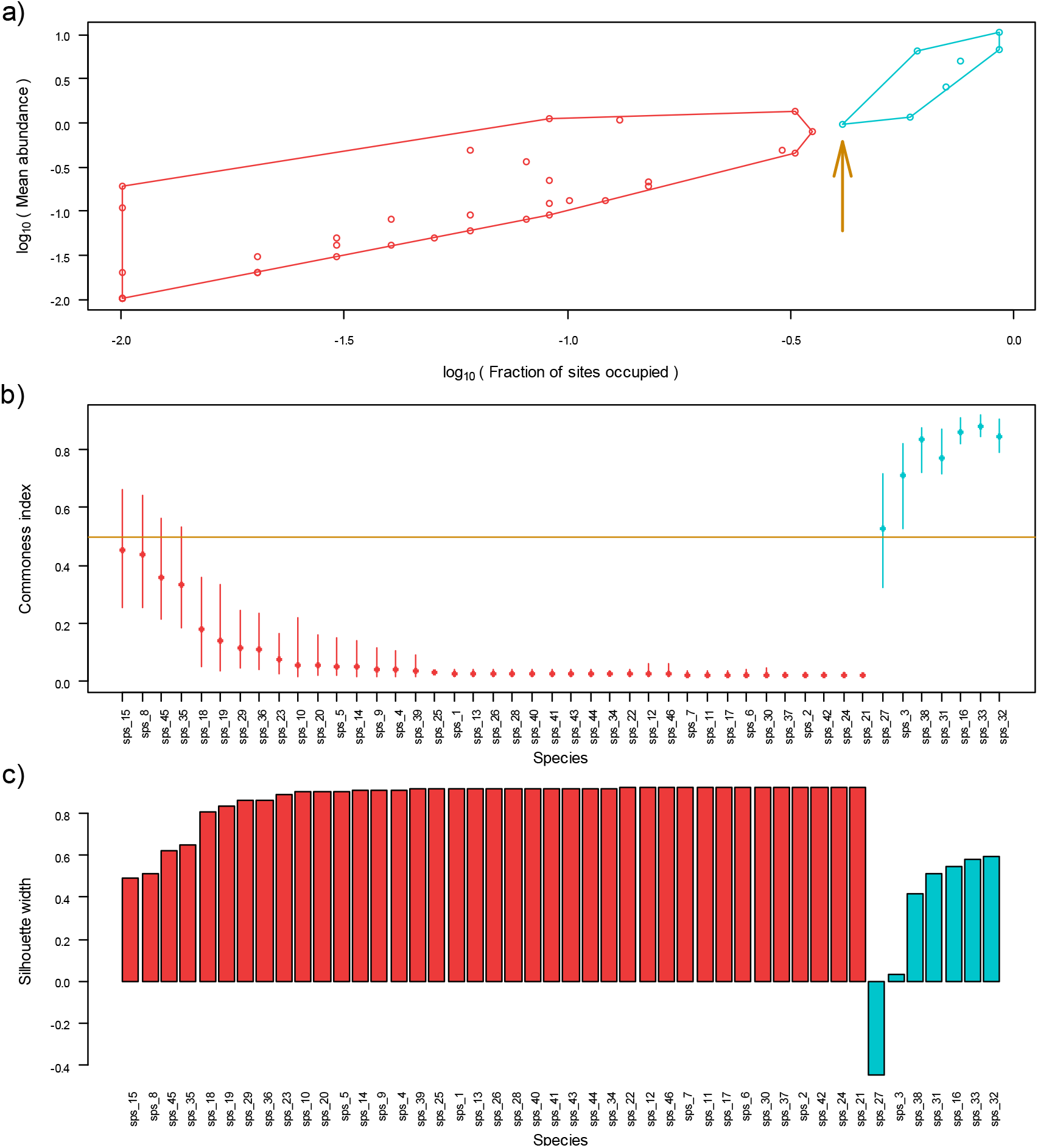
Fuzzy quantification of common and rare species in a community of 46 ant species in 100 plots (Arnan et al., 2011, Calatayud el al., 2019). (a) Abundance occupancy relationship of species. The arrow points to the position of Species 27. (b) Commonness indices of species. Error bars represent bias-corrected and accelerated 95% confidence intervals (error bars) (Efron & Tibshirani, 1994) computed with 1,000 replicates bootstrapping the plots of the abundance matrix. The horizontal line marks the 0.5 threshold separating rare and common species. (c) Silhouette plot of the 46 ant species. The negative value of Species 27 suggests a poor fit to the cluster of common species. Light blue and light red denote common and rare species, respectively.

**TABLE 1.** Overview of the functions implemented in package FuzzyQ.

| Function | Description | Output |
| --- | --- | --- |
| `fuzzyq` | Determines the abundance-occupancy per species of a site × species matrix. Performs fuzzy clustering of common and rare species based on abundance occupancy. | An object of class `list` and `fuzzyq` of three objects:<br>`$A_O`<br>Fraction of sites occupied and the mean abundance across sites per species<br>`$Diss`<br>Object of class `dist` with pairwise dissimilarities among species based on their abundance and occupancy<br>`$spp`<br>Three metrics for each species:<br>1. Cluster membership, where 0 and 1 denote allocation to the rare or common category, respectively<br>2. Silhouette width<br>3. Commonness index<br>`$global`<br>Community-level metrics:<br>1. Average silhouette widths per cluster and globally<br>2. Mean commonness indices per cluster<br>3. Normalized Dunn coefficient |
| `fuzzyqBoot` | Produces $N$ replicates bootstrapping the site × species matrix by site and applies `fuzzyq` to each replicate. | A matrix of either species commonness indices (level = "spp") or community-level metrics (level = "global") of each bootstrap replicate |
| `fuzzyqCI` | Computes confidence intervals of the parameters computed with `fuzzyqBoot`. Three methods are available: percentile, bias corrected, and bias corrected and accelerated. | A matrix of lower and upper bound limits at a given confidence level (default 95%) |
| `AOplot` | Plots the abundance occupancy relationship of a `fuzzyq` object. | A scatter plot of fraction of sites occupied by each species vs mean abundance per site. Common and rare species are distinguished by respective convex hulls |
| `sortClus` | Sorts species data of a matrix (columns) by the cluster allocation of a `fuzzyq` object. Useful to sort the output of `fuzzyqBoot` or `fuzzyqCI` by the original `fuzzyq` object. | A matrix sorted by cluster matrix. Species are arranged by cluster and by increasing silhouette width within cluster |

In fuzzy clustering, each observation can be assigned to several clusters with a different level of certainty. So *u*_*iv*_ in (1) above represents the probability of the *i*^th^ observation belonging to cluster *v* (Kaufman & Rousseeuw, 1990). We re-interpret these probabilities as indices of commonness (*C*_*i*_) and rarity (*R*_*i*_) for species *i*, so that each species is classified simultaneously as common and rare with a certain level of certainty. (Given that *C*_*i*_ = 1 – *R*_*i*_, we will only report *C*_*i*_). `fuzzyqBoot` generates and applies `fuzzyq` to bootstrap replicates by site of the species abundance matrix and `fuzzyqCI` computes confidence intervals of *C*_*i*_ based on these replicates (Fig. 1b).

In addition, `fuzzyq` computes silhouette widths, which are measures of how similar abundance and occupancy of each species are to its own cluster relatives and to these of species in the other cluster, as follows:

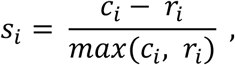

where *ci* and *ri* are the mean dissimilarity between species *i* and all other species in the clusters of common and rare species, respectively (Kaufman & Rousseeuw, 1990). Fig. 1c shows the species silhouettes of the ant database. Silhouettes can range between −1 and +1. The high positive values of most rare ant species indicate that they are well matched to its own cluster. Common ant species showed smaller silhouette widths, suggesting a weaker cluster. In particular, the negative silhouette of species 27 indicates a poor fit to the common-species group (Fig. 1c), which conforms to its position in the AOR plot and its *C*_*i*_ ≈ 0.5 (Fig. 1a, b).

`fuzzyq` also computes community-level metrics that measure the coherence of the common- and rare-species clusters, and the strength of overall classification. The former is assessed by the average silhouettes’ widths of the common and rare species (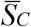 and 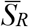, respectively) and, alternatively, by the corresponding average commonness coefficients (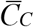 and 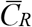). The latter can be appraised by the average silhouette width of the whole community 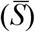 and the normalized Dunn’s partition coefficient (*D’*) (Kaufman & Rousseeuw, 1990). The Dunn’s coefficient is computed as

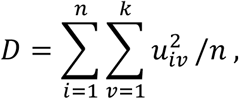

where *n* is the number of observations (i.e., species). *D* is subsequently normalized to vary between 0 (complete fuzziness) and 1 (hard clusters). When *k* = 2, as in our case, the normalized Dunn’s coefficient is *D*′ = 2*D* − 1 (Kaufman & Rousseeuw, 1990).

Fig. 2 displays the global metrics of the ant species database. 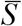 and *D’* were 0.79 and 0.69, which suggests a strong clustering structure separating common and rare species (Kaufman & Rousseeuw, 1990). We assessed the variation of the global estimates by bootstrapping the sites of the sites × species matrix with `fuzzyqBoot`. Compared with common ones, rare species showed a higher average silhouette width, and showed lower variation in both silhouettes and commonness indices, indicating that they form a harder cluster (Fig. 2).

**FIGURE 2.**
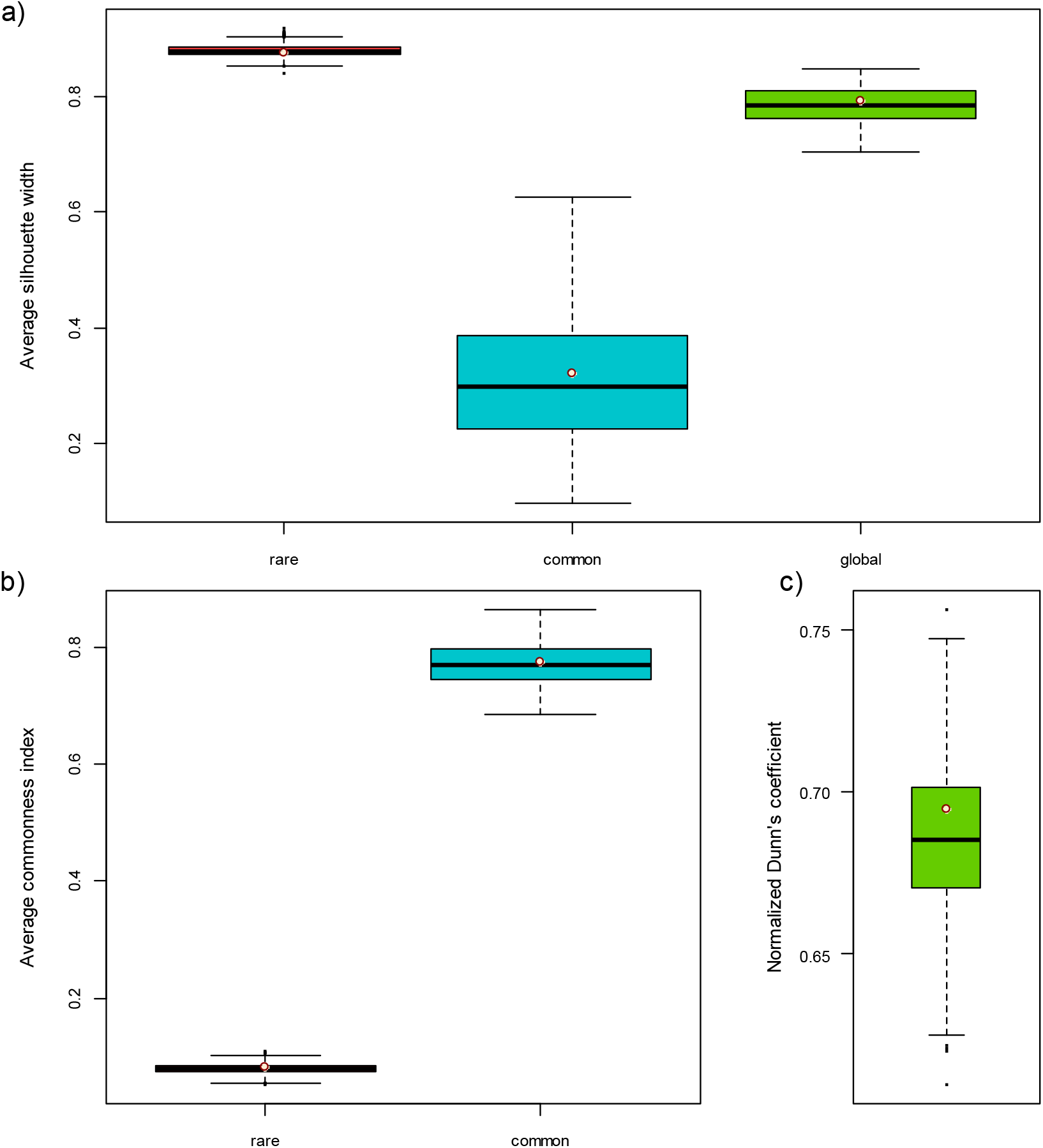
Community-level estimates (points) and their variation computed with 1,000 replicates bootstrapping the plots of the abundance matrix (boxplots) of 46 ant species in 100 plots (Arnan et al., 2011, Calatayud el al., 2019). (a) Average silhouette widths of rare, common and all species. (b) Average commonness indices of rare and common species. (c) Normalized Dunn’s coefficient.

## Worked-out examples

We demonstrate the new method and its capabilities, with two real datasets involving the comparison of related and unrelated communities, respectively. When comparing several communities, one must consider how to deal with species absences. Absences may be due to (a) eco-evolutionary constraints (structural absence), (b) sampling variability (random absence), or (c) methodological errors (false absences) (Blasco-Moreno, Pérez-Casany, Puig, Morante, & Castells, 2019). Although `fuzzyq` cannot deal with (c), it would produce different metrics in (a) and (b) situations and researchers should make an informed decision based on the nature of their system. The logical argument `rm.absent` in `fuzzyq` specifies whether species absences are to be treated as structural or random.

### Example 1. Mammal Data from Powdermill Biological Station 1979-1999

To illustrate how to monitor changes in species commonness in a community, we used a long-term (1979-1999) time series of small mammal abundances from the Powdermill Biological Station in Pennsylvania, USA (Merrit, 2013). Mammals were captured in a 1-ha live trapping grid consisting of 10 × 10 quadrats of trap stations at 10-m intervals. Trapping was conducted bimonthly from September 1979 to October 1999. For the sake of demonstration, the abundance of each mammal species was aggregated per quadrat and per year in order to capture the annual variation in commonness of each species and we assumed that the pool of species did not change over the study period (random absences).

The *C*_*i*_s indicated that two and seven of the 14 species could be categorized consistently as common and rare, respectively, throughout the study period. The *C*_*i*_s of the remaining five varied considerably over the years (Fig. 3a). We modelled the change in commonness of a species in the latter group, *Glaucomys volans* (GV), by fitting a Generalized Additive Model. Bootstrap replicates to fit 95% confidence intervals to the model were generated with `fuzzyqBoot`. The fitted model was also used to predict the change in *C*_*i*_ five years ahead. (Details on model fitting are given in an accompanying R script. See Data Availability below.) The model suggests a progressive increase in *C*_*i*_ of *G. volans* over the years and predicts a similar increase rate in the following years (Fig. 3b).

**FIGURE 3.**
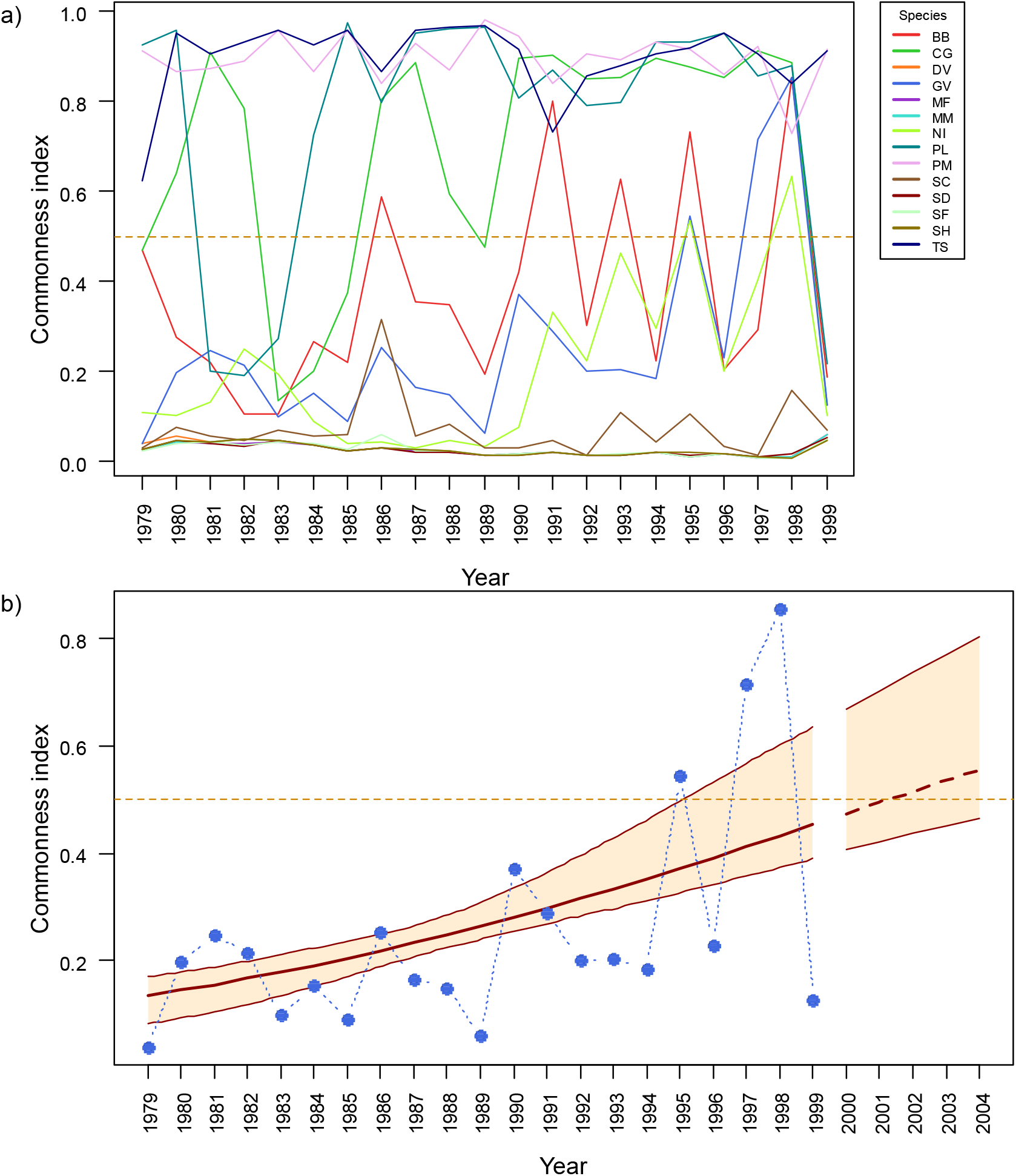
Fuzzy quantification of common and rare species in a community of 14 mammal species sampled at the Powdermill Biological Station from 1979 to 1999 (Merrit, 2013). (a) Variation of commonness indices in the study period. (b) Generalized Additive Model describing the variation in commonness of *Glaucomys volans* (GV) in the study period and predicted change (2001-2004). Blue points: observed values. Thick red line: fitted and predicted model (continuous and stippled lines). Thin red line: 95% confidence interval of the model. Stippled orange line: 0.5 threshold between rare and common species. Species abbreviations: **BB**, *Blarina brevicauda*; **CG**, *Clethrionomys gapperi*; **DV**, *Didelphis virginiana*; **GV**, *Glaucomys volans*; **MF**, *Mustela frenata*; **MM**, *Marmota monax*; **NI**, *Napaeozapus insignis*; **PL**, *Peromyscus leucopus*; **PM**, *Peromyscus maniculatus*; **SC**, *Sorex cinereus*; **SD**, *Sorex dispar*; **SF**, *Sorex fumeus*; **SH**, *Sorex hoyi*; **TS**, *Tamias striatus*.

### Example 2. Parasite communities of the so-iuy mullet in native and introduced areas

We compared the patterns of commonness and rarity of helminth communities of the so-iuy mullet (*Planiliza haematocheilus* in its native (Sea of Japan) and introduced (Sea of Azov and Black Sea) areas (Llopis-Belenguer, Blasco-Costa, Balbuena, Sarabeev, & Stouffer, 2020). We used here 12 and 7 surveys in the introduced and native areas, respectively, in which the number of fish sampled was ≥ 20, totalling 378 and 192 fish, respectively. Based on biogeographical evidence, species absences within the native and introduced areas were treated as random zeros (Kostadinova, 2008).

We used `fuzzyq` to compute 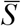, 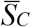, 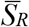, 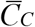, 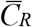 and *D’* of each survey and `fuzzyqBS` and `fuzzyqCI` to estimate their 95% confidence intervals (Fig. 4). Differences in these metrics between surveys in the native and introduced areas were evaluated by Mann-Whitney tests. In the introduced area, rare species had significantly higher 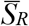 and lower 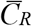 than in the native one (*p* = 0.0012 and *p* = 0.0003, respectively). Differences in 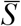 and *D’* were also significant (*p* = 0.0003 and *p* = 0.0002, respectively), indicating a clearer distinction between common and rare species in the introduced area than in the native one. By contrast, there was no evidence for significant differences between areas in 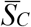 and 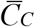 (*p* = 0.71 and *p* = 0.97, respectively). These results conform to previous work that indicates that the introduction of the mullet so-iuy in the new area entailed a deep structural change in its helminth communities (Sarabeev, Balbuena, & Morand, 2017; Llopis-Belenguer et al., 2020). Most native species were lost and only two *Ligophorus* spp. common in the native area were co-introduced and remained common in the introduced area (Figs. S1, S2 in Supporting Information). So the majority of species in the introduced area were acquired from local grey mullet species (Sarabeev et al., 2017). Since newly acquired parasite species are expected to lack specific adaptations to the new host, this would account for their pronounced rarity compared to rare species in the native area (Sarabeev, Balbuena, & Morand, 2018).

**FIGURE 4.**
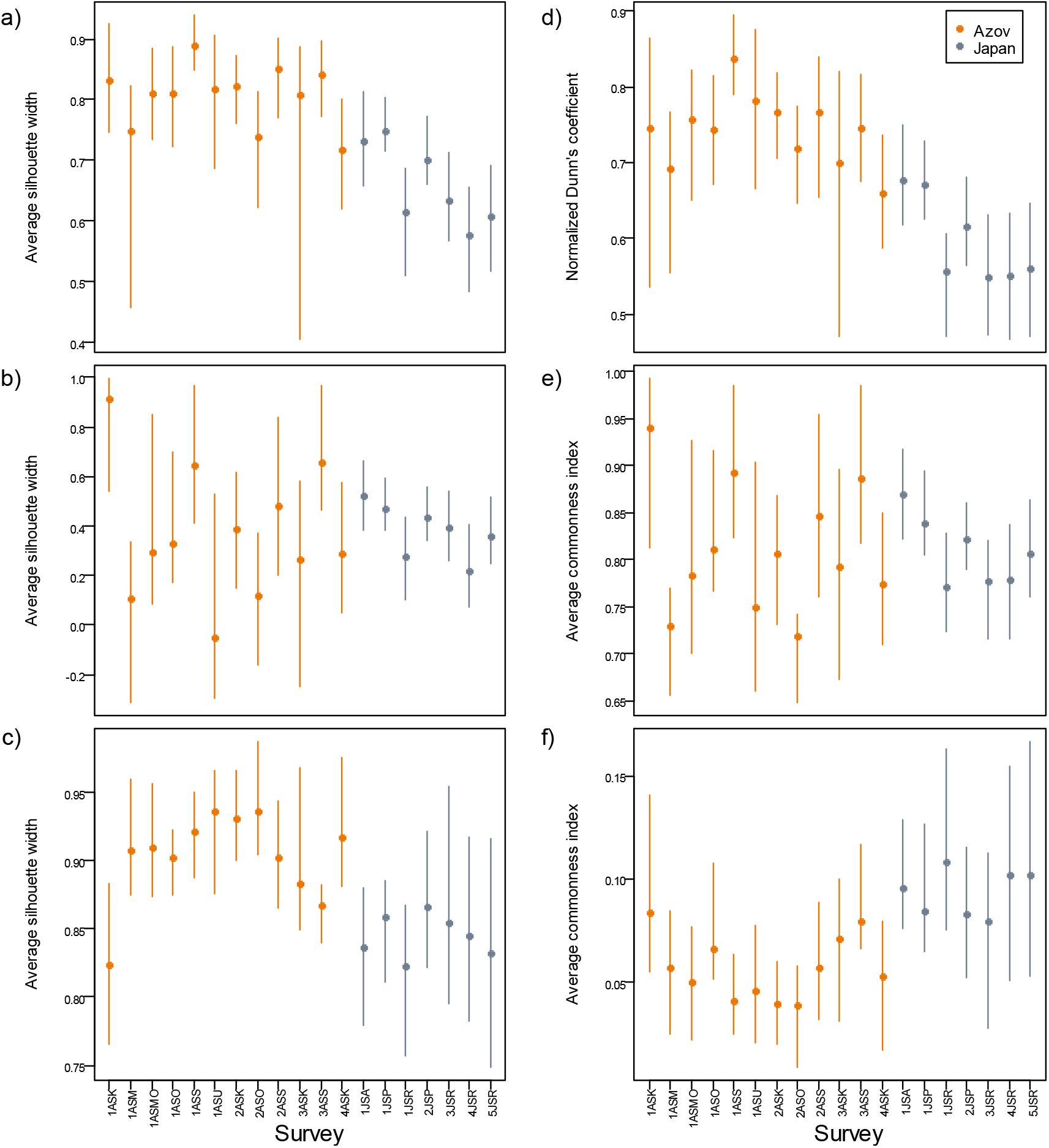
Community-level estimates (points) and bias-corrected and accelerated 95% (Efron & Tibshirani, 1994) confidence intervals (error bars) computed with 1,000 replicates bootstrapping the plots of the abundance matrix of helminth communities of *Planiliza haematocheilus* in 7 native (Japan Sea) and 12 introduced (Azov and Black Seas) surveys. (a) Average silhouette widths of all species. (b) Idem common species. (c) Idem rare species. (d) Normalized Dunn’s coefficient. (e) Average commonness indices of common species. (f) Idem rare species.

## Effect of number of sites

A key question for users interested in comparing different communities is whether the community-level estimates depend on the number of sites sampled. We examined this issue using 20 datasets compiled in Calatayud et al. (2019) and (Jeliazkov et al., 2020) involving 87+ sites. These included communities varying widely in taxonomic composition, species richness and spatial scale. In addition, different abundance currencies were employed. (See Table S1 in Supporting Information for details.) Being *N* the total number of sites of a given dataset, global metrics (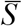, 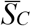, 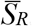, 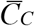, 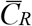 and *D’*) were computed in most cases dropping successively 1, 2, 3, …, *N*-10 sites randomly drawn (without replacement) from the dataset. (In species-poor communities or communities with very sparsely distributed species, the series was 1, 2, 3, …, *N*-20). Species absences in each draw were treated as structural, because our goal was to examine how incomplete species coverage resulting from low sample sizes affected the estimation of global metrics.

Fig. 5 shows the variation of the global metrics with the number of sites in eight of the 20 datasets. (Results for the remaining 12 datasets are given in Fig. S3, Supporting Information.) No clear trend of variation with the number of sites was apparent (Fig. 5 and Fig. S3, Supporting Information). Although in some datasets large fluctuations occurred (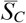 in particular was quite labile in some examples), variation in global parameters did not seem related to sample size. The results suggest that 30 to 50 sites are sufficient to yield reliable estimates although bootstrapping should be used to capture their variability.

**FIGURE 5.**
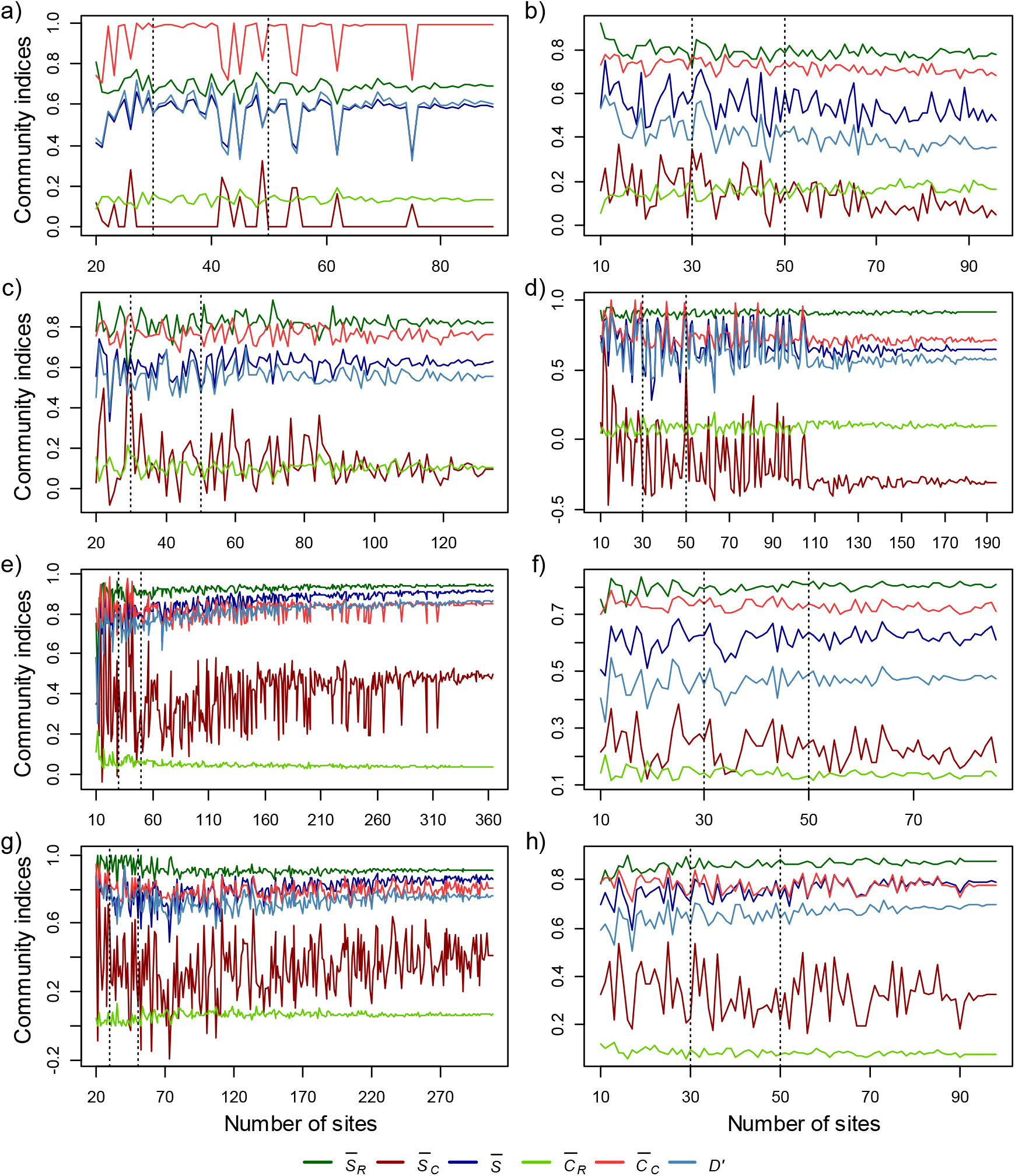
Variation of community-level metrics with number of sites in eight, 87+-site, databases from Jeliazkov et al. (2020) (a-g) and Calatayud et al. (2019) (h): (a) BrindAmour2011a; (b) Pavoine2011; (c) Jeliazkov2014; (d) Barbaro2009a; (e) Chmura2016; (f) Ribera2001; (g) Goncalves2014a; (h) ants_data_Xavi_Darwin_A. Abbreviations: 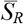, average silhouette rare species; 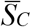, idem common species; 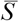, idem all species; 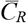, Commonness coefficient rare species; 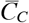, idem common species; *D’*, Normalized Dunn’s coefficient. Details of these datasets are given in Table S1, Supporting Information.

## Final remarks

FuzzyQ provides a new quantitative framework to study the distribution of common and rare species in ecological communities. The approach supplies simple and intuitive ecological indicators that can give both clear, actionable insights into the nature of ecological communities and a powerful way to monitor quantitatively environmental change on ecosystems. We show that the approach works satisfactorily with a wide range of communities varying in species richness, dispersion and abundance currencies. The only obvious limitation in its application is that fuzzy clustering requires that *k* ≥ *n*/2 −1 (Kaufman & Rousseeuw, 1990). As in our case *k* = 2, FuzzyQ cannot be applied to communities composed of ≤ 5 species. In addition, the application of fuzzyqBoot in communities with low number of species can lead to a number of null replicates because of this limitation.

Comparison among communities is greatly facilitated by the fact that the method is relatively independent of the number of sites or sampling units considered. However, the use of FuzzyQ in comparative settings comes with an important caveat. Since FuzzyQ is based on AORs and occupancy is known to vary with spatial scale (Hui, Veldtman, & McGeoch, 2010; Steenweg, Hebblewhite, Whittington, Lukacs, & McKelvey, 2018), differences in scaling can compromise comparison among communities. In our second working example, helminth communities of individual fish were evaluated as sites. Therefore, we consider that the comparison makes biological sense. However, we cannot completely rule out that potential differences between the native and introduced areas in fish mobility could introduce a hidden bias (Steenweg et al., 2018).

Likewise, it has been shown that rarity at coarse scales can be substantially biased because species of similar occupancies at that level may have very different occupancies at finer scales (He & Condit, 2007). Thus, assessment and monitoring of rarity should be performed at the appropriate scale for suitable conservation and management plans. Nevertheless, for samples taken at nested spatial or temporal scales, FuzzyQ provides a convenient tool to assess how scale affects patterns of commonness and rarity. In addition, the approach is versatile as it can be readily adapted to other categorizations (by considering more clusters) or to other criteria of rarity (by introducing additional/different traits when computing the dissimilarity matrix). Therefore, we consider that FuzzyQ is a potentially valuable analytical tool in community ecology and conservation biology.

## Data Availability

Table 2 provides information of the availability of the datasets used herein. R scripts and R markdown files to run the illustrative examples, and the FuzzyQ R package are available at https://ligophorus.github.io/FuzzyQ/ (**DOI no. pending**).

**TABLE 2.**
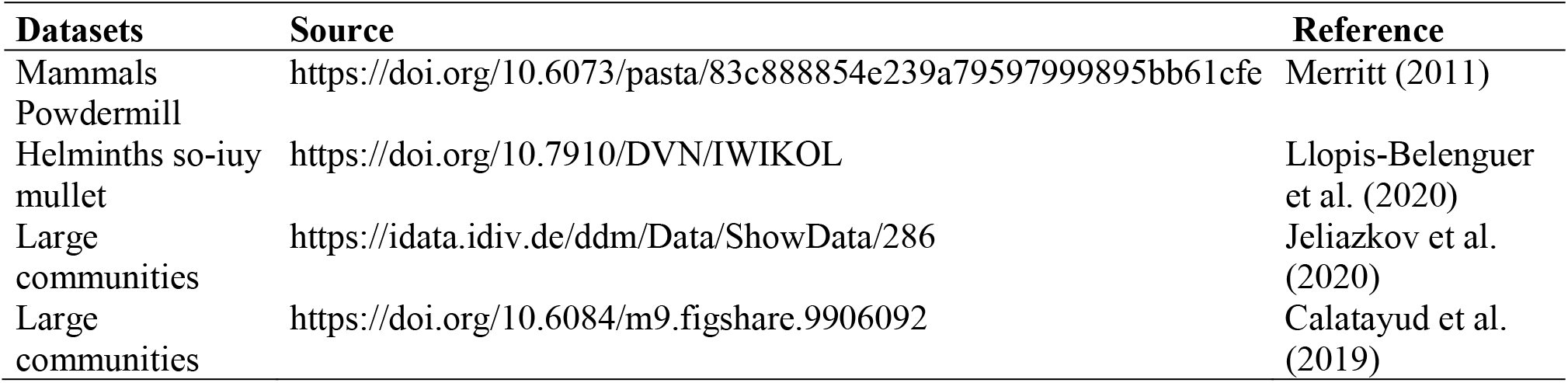
Availability of the datasets used with FuzzyQ.

## Acknowledgements

We thank Xavier Arman and co-workers for allowing us to incorporate their ant species dataset to the FuzzyQ package. Study funded by the Ministry of Science and Innovation, Spain (PID2019-104908GB-I00). The mammal Powdermill data was obtained with the support of NSF Grants BSR-8702333-06, DEB-9211772, DEB-9411974, DEB-0080381 and DEB-0621014.

## Authors’ contributions

JAB conceived the idea. JAB and SM developed the theory and outlined the study. CM, CLB and JAB wrote the scripts and developed the package. CLB, IBC, VLS and SM set and verified the analytical methods. All authors contributed decisively to shape the research, provided critical feedback on the drafts and gave final approval for submission.

## Notes

https://ligophorus.github.io/FuzzyQ/

### Competing Interest Statement

The authors have declared no competing interest.

