## Supplementary Information for "Fuzzy Quantification of Common and Rare Species in Ecological Communities (FuzzyQ)"

This document provides supporting tables and figures to the accompanying paper.

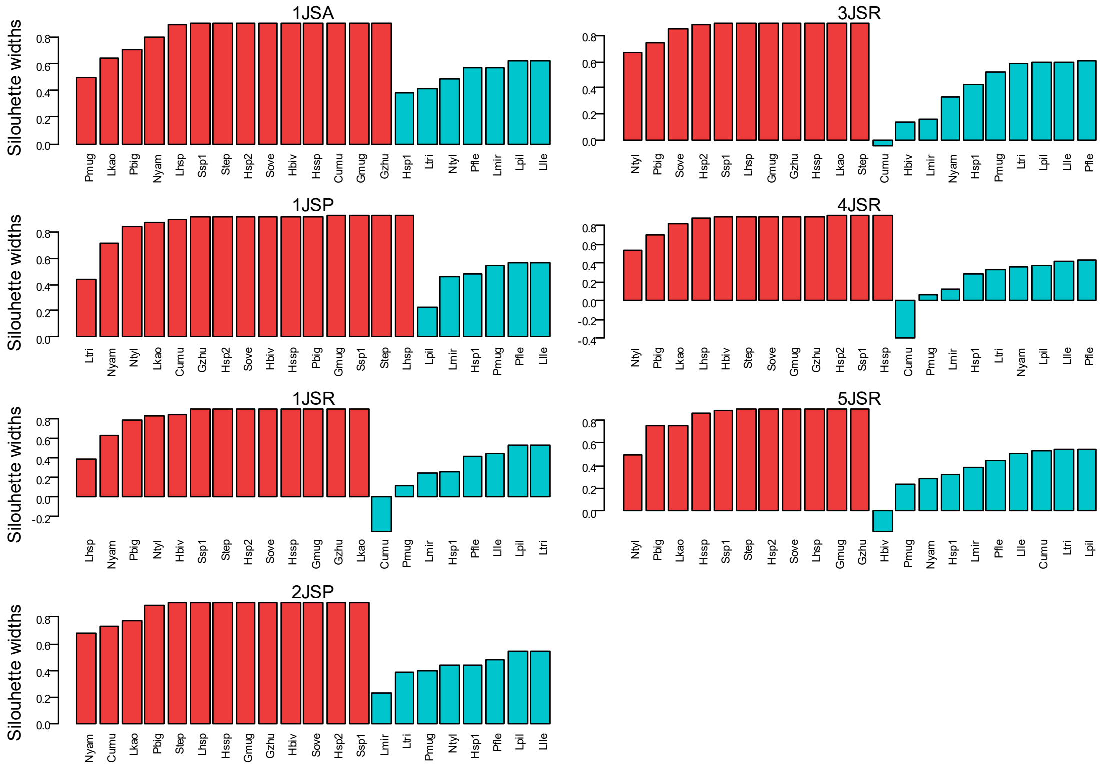


**Figure S1.** Silhouette plots of helminth species of the so-iuy mullet *Planiliza haematocheilus* from 7 surveys in the Japan Sea (native area). Light blue and light red denote common and rare species, respectively. Survey ID and species abbreviations in Llopis-Belenguer (2019).

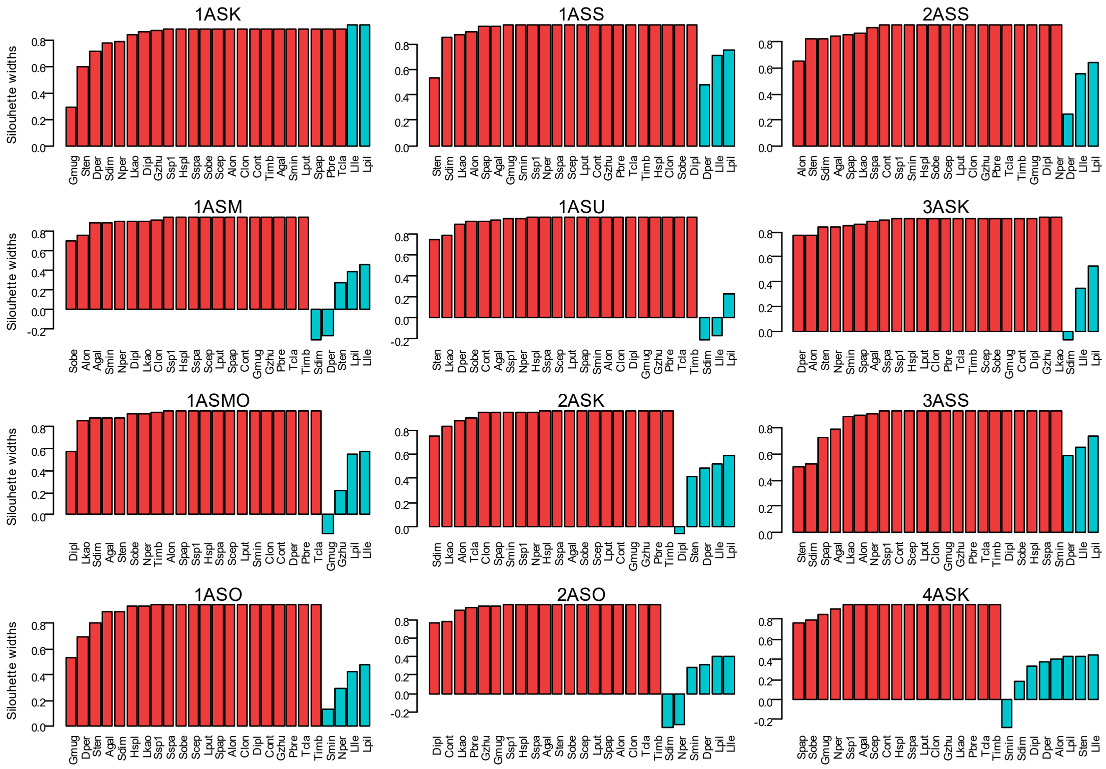


**Figure S2.** Silhouette plots of helminth species of the so-iuy mullet *Planiliza haematocheilus* from 12 surveys in the Azov and Black Seas (introduced area). Light blue and light red denote common and rare species, respectively. Survey ID and species abbreviations in Llopis-Belenguer (2019).

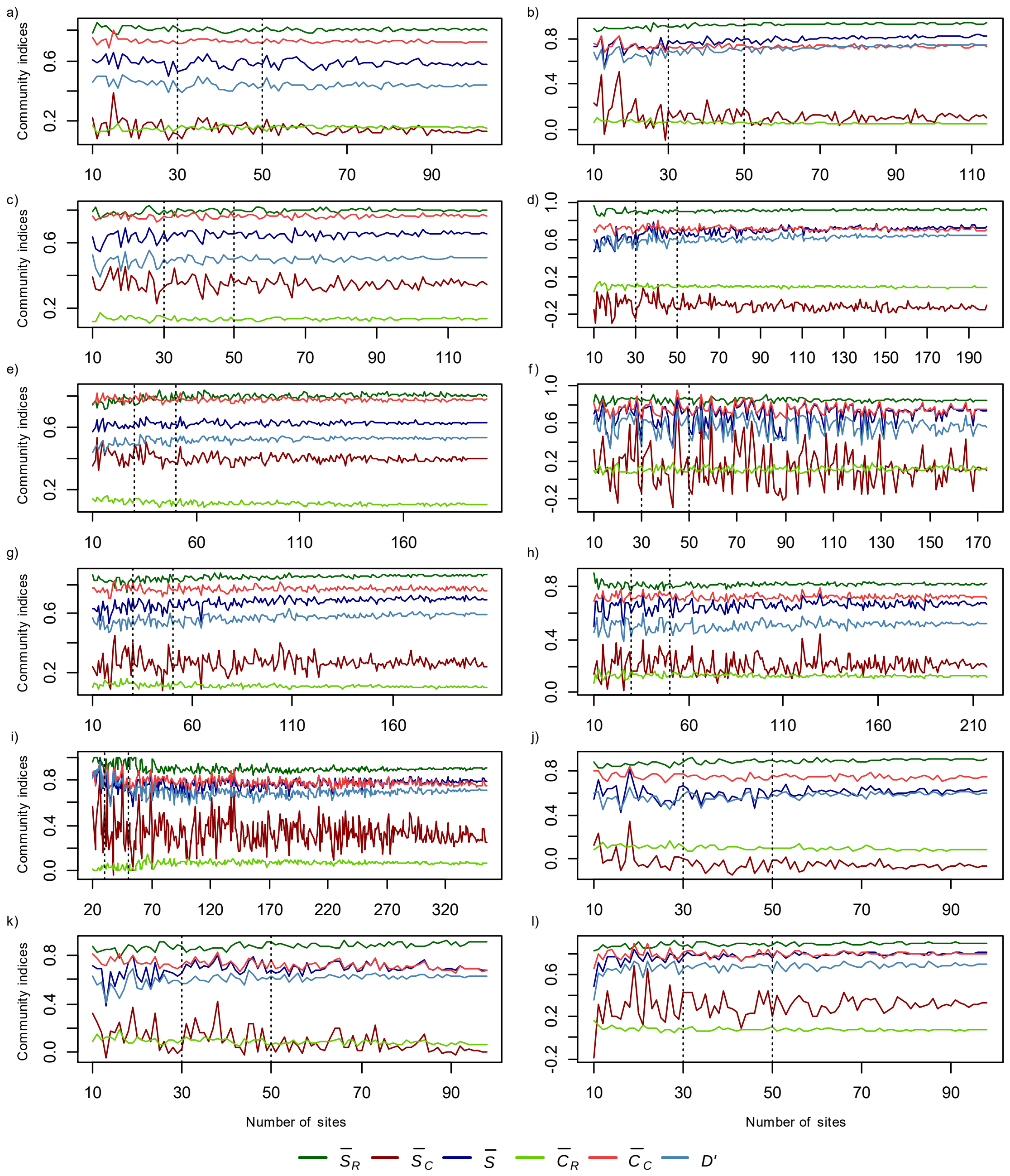


**Figure S3.** Variation of community-level metrics with number of sites in eight, 87+-site, databases from Jeliazkov et al. (2020) (*a-i*) and Calatayud et al. (2019) (*j-l*): (*a*) Diaz2008; (*b*) Lowe2018a; (*c*) Bartonova2016; (*d*) Jeliazkov2013; (*e*) Barbaro2009b; (*f*) Charbonier2016a; (*g*) Charbonier2016b; (*h*) Fried2012; (*i*) Goncalves2014b; (*j*) ants_data_Xavi_Darwin_B; (*k*) ants_data_Xavi_Darwin_C; (*l*) ants_data_Xavi_Darwin_D. Abbreviations: $\bar{S}_R$, average silhouette rare species; $\bar{S}_C$, idem common species; $\bar{S}$, idem all species; $\bar{C}_R$, Commonness coefficient rare species; $\bar{C}_C$, idem common species; *D'*, Normalized Dunn's coefficient. Details of these datasets are given in Table S1.

**Table S1.** Information on 20 datasets compiled involving 87+ sites used to assess the effect of sample size on the community-level metrics computed by FuzzyQ.

| Dataset name | Source[a] | Type of data[b] | Taxonomic group | Ecosystem type | Extent ($km^2$) | N spp.[c] | N sites[d] | Reference |
|---|---|---|---|---|---|---|---|---|
| ants_data_Xavi_Darwin_A | C | NI | Ants | Terrestrial | 0.00032 | 46 | 100 | Arnan et al. (2011) |
| ants_data_Xavi_Darwin_B | C | NI | Ants | Terrestrial | 0.00032 | 38 | 100 | Arnan et al. (2011) |
| ants_data_Xavi_Darwin_C | C | NI | Ants | Terrestrial | 0.00032 | 40 | 100 | Arnan et al. (2011) |
| ants_data_Xavi_Darwin_D | C | NI | Ants | Terrestrial | 0.00032 | 37 | 100 | Arnan et al. (2011) |
| Barbaro2009a | J | NI | Beetles | Terrestrial | 32.16 | 36 | 195 | Barbaro & van Halder (2009) |
| Barbaro2009b | J | AI | Birds | Terrestrial | 32.16 | 53 | 201 | Barbaro & van Halder (2009) |
| Bartonova2016 | J | AC | Butterflies | Terrestrial | 78866 | 128 | 122 | Bartonova et al. (2016) |
| BrindAmour2011a | J | AC | Fishes | Freshwater | 0.31 | 7 | 90 | Brind'Amour et al. (2011) |
| Charbonier2016a | J | NI | Bats | Terrestrial | 4400000 | 27 | 175 | Charbonier et al. (2016) |
| Charbonier2016b | J | NI | Birds | Terrestrial | 4400000 | 73 | 208 | Charbonier et al. (2016) |
| Chmura2016 | J | NI | Plants | Terrestrial | 135.05 | 46 | 364 | Chmura et al. (2016) |
| Diaz2008 | J | NI | Macroinvertebrates | Freshwater | 6300 | 208 | 104 | Mellado-Díaz et al. (2008) |
| Fried2012 | J | DE | Plants | Terrestrial | 386000 | 75 | 218 | Fried et al. (2012) |
| Goncalves2014a | J | NI | Spiders | Terrestrial | 220000 | 105 | 309 | Gonçalves-Sousa et al. (2014) |
| Goncalves2014b | J | NI | Arthorpods | Terrestrial | 220000 | 112 | 356 | Gonçalves-Sousa et al. (2014) |
| Jeliazkov2013 | J | NI | Macroinvertebrates | Freshwater | 430 | 91 | 112 | Jeliazkov (2013) |
| Jeliazkov2014 | J | NI | Amphibians | Freshwater | 430 | 11 | 135 | Jeliazkov et al. (2014) |
| Lowe2018a | J | NI | Spiders | Terrestrial | 1000 | 135 | 115 | Lowe et al. (2018) |
| Pavoine2011 | J | NI | Plants | Terrestrial | 100 | 56 | 97 | Pavoine et al. (2011) |
| Ribera2001 | J | NI | Beetles | Terrestrial | 78772 | 68 | 87 | Ribera et al. (2001) |

[a] C: Calatayud et al. (2019); J: Jeliazkov et al. (2020). [b] AC: Abundance class; AI: Abundance Index; DE: Density (individuals/$m^2$); NI: Number of Individuals. [c] N spp.: Number of species. [d] N sites: Number of sites
